# Chemical reprogramming ameliorates cellular hallmarks of aging and extends lifespan

**DOI:** 10.1101/2022.08.29.505222

**Authors:** Lucas Schoenfeldt, Patrick T. Paine, Nibrasul H. Kamaludeen M., Grace B. Phelps, Calida Mrabti, Kevin Perez, Alejandro Ocampo

**Affiliations:** Department of Biomedical Sciences, Faculty of Biology and Medicine, University of Lausanne, Lausanne, Vaud, Switzerland

**Author notes:** These authors contributed equally (Lucas Schoenfeldt, Patrick T. Paine).

**Keywords:** Aging, cellular reprogramming, chemical reprogramming, epigenetics, lifespan

## Abstract

The dedifferentiation of somatic cells into a pluripotent state by cellular reprogramming coincides with a reversal of age-associated molecular hallmarks. Although transcription factor induced cellular reprogramming has been shown to ameliorate these aging phenotypes in human cells and extend health and lifespan in mice, translational applications of this approach are still limited. More recently, chemical reprogramming via small molecule cocktails have demonstrated a similar ability to induce pluripotency in vitro, however, its potential impact on aging is unknown. Here, we demonstrated that partial chemical reprogramming is able to improve key drivers of aging including genomic instability and epigenetic alterations in aged human cells. Moreover, we identified an optimized combination of two reprogramming molecules sufficient to induce the amelioration of additional aging phenotypes including cellular senescence and oxidative stress. Importantly, in vivo application of this two-chemical combination significantly extended *C. elegans* lifespan. Together, these data demonstrate that improvement of key drivers of aging and lifespan extension is possible via chemical induced partial reprogramming, opening a path towards future translational applications.

## INTRODUCTION

Biological aging is a global process associated with a loss of homeostasis and functional decline across cellular and physiological systems leading to the development of age-associated chronic diseases and finally death (Rando and Chang, 2012; Kennedy *et al*., 2014). For this reason, a significant increase in human life expectancy over the last decades has resulted in an extended period of life spent in morbidity (Garmany, Yamada and Terzic, 2021). As aging is a key risk factor for chronic diseases such as cardiovascular disease or neurodegenerative disorders, new therapeutic strategies that target aging are now under intense investigation (Niccoli and Partridge, 2012; Mahmoudi, Xu and Brunet, 2019). Several hallmarks of aging including epigenetic dysregulation, genomic instability, cellular senescence, and stem cell exhaustion have been identified as potential targets for such an optimized therapeutic strategy (López-Otín *et al*., 2013). Current interventions that target aging include cellular reprogramming, dietary restriction and related mimetics, systemic blood factors, and senolytics. Among these, cellular reprogramming offers a unique prospective for its ability to reset the epigenome and restore multiple aging hallmarks (Conboy *et al*., 2005; Baker *et al*., 2011; Lapasset *et al*., 2011; Longo *et al*., 2015; Ocampo *et al*., 2016; Olova *et al*., 2019).

During development, cellular reprogramming induces zygotic and primordial germ cell formation following a dramatic chromatin reorganization to create totipotent and pluripotent cells free of aged molecular defects demonstrating that both cell identity and age are reversible (Seisenberger *et al*., 2013; Kerepesi *et al*., 2021). Importantly, this manipulation of cell identity has been recapitulated in vitro by several methods including somatic cell nuclear transfer, forced expression of transcription factors, and most recently treatment with small molecules (Gurdon, 1962; Takahashi and Yamanaka, 2006; Hou *et al*., 2013).

Although restoration of aged phenotypes such as telomere length, mitochondrial function, proliferation, and transcriptomic signature in vitro was demonstrated over a decade ago, application of cellular reprogramming in vivo was initially proven unsafe due to the loss of cellular identity leading to tumor and teratoma formation (Lapasset *et al*., 2011; Abad *et al*., 2013). To overcome this issue, in vivo partial reprogramming by short-term cyclic induction of Oct4, Sox2, Klf4, and c-Myc (OSKM) was a critical advance as it avoided the detrimental loss of cellular identity. Importantly, this limited cyclic expression of OSKM was sufficient to ameliorate multiple aging hallmarks and extend the lifespan of a progeroid mouse strain (Ocampo *et al*., 2016). Improved regenerative capacity and function has also been demonstrated following therapeutic application of cellular reprogramming in multiple tissues and organs including the intervertebral disc, heart, skin, skeletal muscle, liver, optic nerve, and dentate gyrus (Ocampo *et al*., 2016; Kurita *et al*., 2018; de Lázaro *et al*., 2019; Lu *et al*., 2020; Rodríguez-Matellán *et al*., 2020; Chen *et al*., 2021; Cheng *et al*., 2022; Hishida *et al*., 2022). Furthermore, several groups have demonstrated the ability to restore multiple aging phenotypes and reset the epigenetic clock utilizing translational non-integrative methods such as modified mRNAs encoding for six transcription factors (OSKM + Lin28 and Nanog) or adeno-associated virus for expression of three factors (OSK) (Sarkar *et al*., 2020; Lu *et al*., 2020). While promising, methods that require transcription factor expression face significant barriers for their clinical translation such as risk of tumorigenicity and low delivery efficiency (Abad *et al*., 2013; Ohnishi *et al*., 2014). In this line, c-Myc and Klf4 have been identified as proto-oncogenes while Oct4 and Sox2 are highly expressed in a variety of human cancers (Klimczak, 2015). For this reason, the clinical application of in vivo reprogramming may require further development.

Most recently, small molecule cocktails have been shown to produce chemical induced pluripotent stem cells (ciPSCs) from mouse and human somatic cells (Hou *et al*., 2013; Guan *et al*., 2022). These reprogramming compounds fall broadly into three categories including epigenetic, cell signaling, and metabolic modulators (Knyazer *et al*., 2021). Importantly, small molecule reprogramming and OSKM expression share the ability to overcome multiple reprogramming barriers while retaining a distinct cell fate trajectory (Zhao *et al*., 2015; Haridhasapavalan *et al*., 2020). To date, the effects of chemical reprogramming on aging hallmarks and lifespan have not been investigated. Considering both the rejuvenating effects of partial reprogramming by short-term expression of OSKM and the ability of small molecule cocktails to induce pluripotency, we proposed the use of chemical induced partial reprogramming for the amelioration of aging phenotypes.

Here, we report that short-term treatment of human cells with seven small molecules (7c), previously identified for their capacity to induce pluripotent stem cells, leads to the improvement of molecular hallmarks of aging. In addition, we show that an optimized cocktail, containing only two of these small molecules (2c), is sufficient to restore multiple aging phenotypes including genomic instability, epigenetic dysregulation, cellular senescence, and elevated reactive oxygen species. Finally, in vivo application of this 2c reprogramming cocktail extends lifespan in *C. elegans*.

## RESULTS

### Chemical induced partial reprogramming significantly improves aging hallmarks in aged human fibroblasts

Multiple hallmarks of aging can be ameliorated following partial cellular reprogramming by expression of the Yamanaka factors (OSKM) in vitro and in vivo (Ocampo, Reddy and Belmonte, 2016). On the other hand, although chemical reprogramming with seven small molecules (7c) has been shown to generate chemically induced pluripotent stem cells (Hou *et al*., 2013), whether chemical induced partial reprogramming is also able to restore aged phenotypes is unknown. Therefore, we sought to determine the effect of short-term 7c treatment on aging phenotypes in primary aged human fibroblasts (Fig. 1a). Specifically, we asked whether chemical induced partial reprogramming could improve multiple hallmarks of aging, including DNA damage, heterochromatin loss, cellular senescence, and reactive oxygen species (ROS) in vitro. Towards this goal, primary human fibroblasts isolated from aged dermal tissue samples were treated for 6 days with a 7c cocktail including: CHIR99021, DZNep, Forskolin, TTNPB, Valproic acid (VPA), Repsox, and Tranylcypromine (TCP). Notably, the levels of the DNA damage marker γH2AX were significantly decreased in aged cells after treatment (Fig. 1b). Interestingly, a decrease in γH2AX levels was also observed when 7c was added for 6 days to aged cells that were pretreated with the DNA damage inducing agent doxorubicin for 2 days, indicating a significantly improved DNA damage response upon 7c treatment (Fig. 1c). Thus, we determined that short-term 7c treatment improves DNA damage in primary aged human fibroblasts.

**Figure 1.**
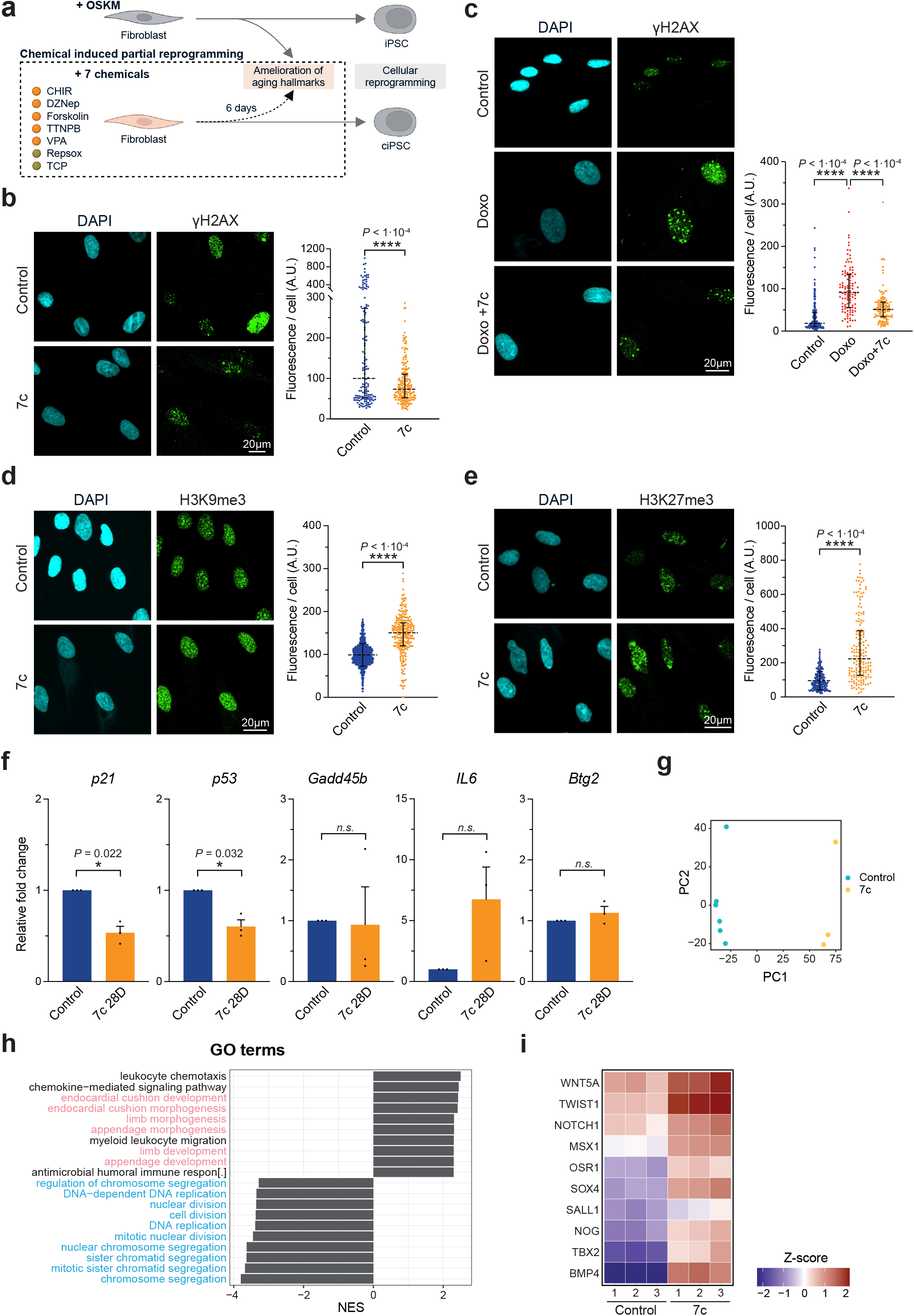
Chemical induced partial reprogramming significantly improves aged hallmarks in aged human fibroblasts. **a**, Schematic representation of chemical induced partial reprogramming via 6 days treatment with the 7 chemicals previously shown to induce mouse chemical iPSC (Hou *et al*., 2013). **b-c**, Immunofluorescence and quantification of γH2AX following 7c treatment (**b**) or Doxorubicin (100 nM) and 7c treatment (**c**). **d-e**, Immnofluorescence and quantification of H3K9me3 (**d**) and H3K27me3 (**e**) following 7c treatment. **f**, mRNA levels of senescence-associated and age-related stress response genes in the *p53* tumor suppressor pathway following 7c treatment during replicative induced senescence (RIS; 28 days). **g**, Principal component analysis (PCA) of control (blue) and 7c treated (orange) fibroblasts. **h**, Gene ontology (GO) enrichment analysis following 7c treatment with developmental (in pink) and cell cycle (in blue) pathways highlighted. **i**, List of differentially expressed genes associated with developmental pathways following 7c treatment. Data are median ± IQR (**b-c**), mean ± SEM (**d-f**). (**b-c**) n=2, (**d-i**) n=3. Statistical significance was assessed by comparison to untreated control using paired two-tailed *t*-test (**b-f**). NES, Normalized enrichment scores.

Epigenetic dysregulation and heterochromatin loss are key molecular markers of aging (Haithcock *et al*., 2005; Scaffidi and Misteli, 2006; Ni *et al*., 2012; Brunet and Berger, 2014; Djeghloul *et al*., 2016; Kane and Sinclair, 2019). For this reason, we next examined the effect of 7c treatment on the constitutive and facultative heterochromatin marks H3K9me3 and H3K27me3. Our results show that 6 days of 7c treatment significantly increased the constitutive heterochromatin mark H3K9me3 in aged human fibroblasts (Fig. 1d). In aged cells, previous work has shown that the facultative heterochromatin marker H3K27me3 is decreased at the senescence-associated *p16* gene locus (*CDKN2A*) leading to increased expression, cell cycle arrest, and senescence (Dhawan, Tschen and Bhushan, 2009). Interestingly, we observed that H3K27me3 was significantly increased after 6 days of 7c treatment (Fig. 1e). Next, as cellular senescence has been shown to be a key driver of aged tissue dysfunction and ablation of senescent cells has been demonstrated to extend health and lifespan in mice (Baker *et al*., 2011), we determined the impact of chemical induced partial reprogramming on senescence-associated gene expression. For this purpose, we serially passaged aged human fibroblasts for 28 days to promote replicative induced senescence (RIS) in the presence of continuous 7c treatment. Importantly, the 7c treated group showed a downregulation of the senescence-associated cell cycle genes *p21* and *p53* relative to untreated controls (Fig. 1f). On the other hand, we noted that the senescence-associated secretory cytokine *IL6* was significantly upregulated upon long-term treatment with 7c and therefore could not conclude a positive effect of 7c treatment on senescence (Fig. 1f and S1a).

Next, further characterization of the transcriptomic effects of 7c treatment for 6 days was performed by bulk RNA sequencing on aged fibroblasts treated with either 7c or vehicle control. Principle component analysis (PCA) showed that 7c treated cells clustered separately from the control group indicating that a distinct transcriptomic profile emerges following chemical induced partial reprogramming (Fig. 1g). Gene ontology (GO) enrichment analysis revealed that developmental processes were significantly upregulated following 7c treatment relative to control, whereas mitosis and cell proliferation programs were significantly downregulated (Fig. 1h). Interestingly, numerous cellular reprogramming, stem cell, and self-renewal genes within the GO term developmental pathways were significantly upregulated following 7c treatment compared to control including *WNT5a, NOTCH1A, SOX4, SALL1, NOG*, and *BMP4* indicating that 7c induced a shift towards a developmental associated transcriptomic profile (Fig. 1i and S1b).

Since our RNA seq analysis indicated a downregulation in mitosis and proliferation programs, we next evaluated the effect of 7c treatment on cellular proliferation. In agreement with these results, we observed that 7c significantly decreased cell density based on MTS assay (Fig. 2a). This observation was confirmed by a strong decrease in the proliferation associated marker Ki67 in cells treated with 7c (Fig. 2b). Subsequently, to determine whether the effect of 7c on proliferation was dose-dependent, a serial dilution assay of 7c treatment was performed. Notably, different concentrations of 7c continued to impair proliferation (Fig. S2a). Overall, these results validate our transcriptomic findings indicating that mitosis and proliferation related programs are downregulated upon 7c treatment.

**Figure 2.**
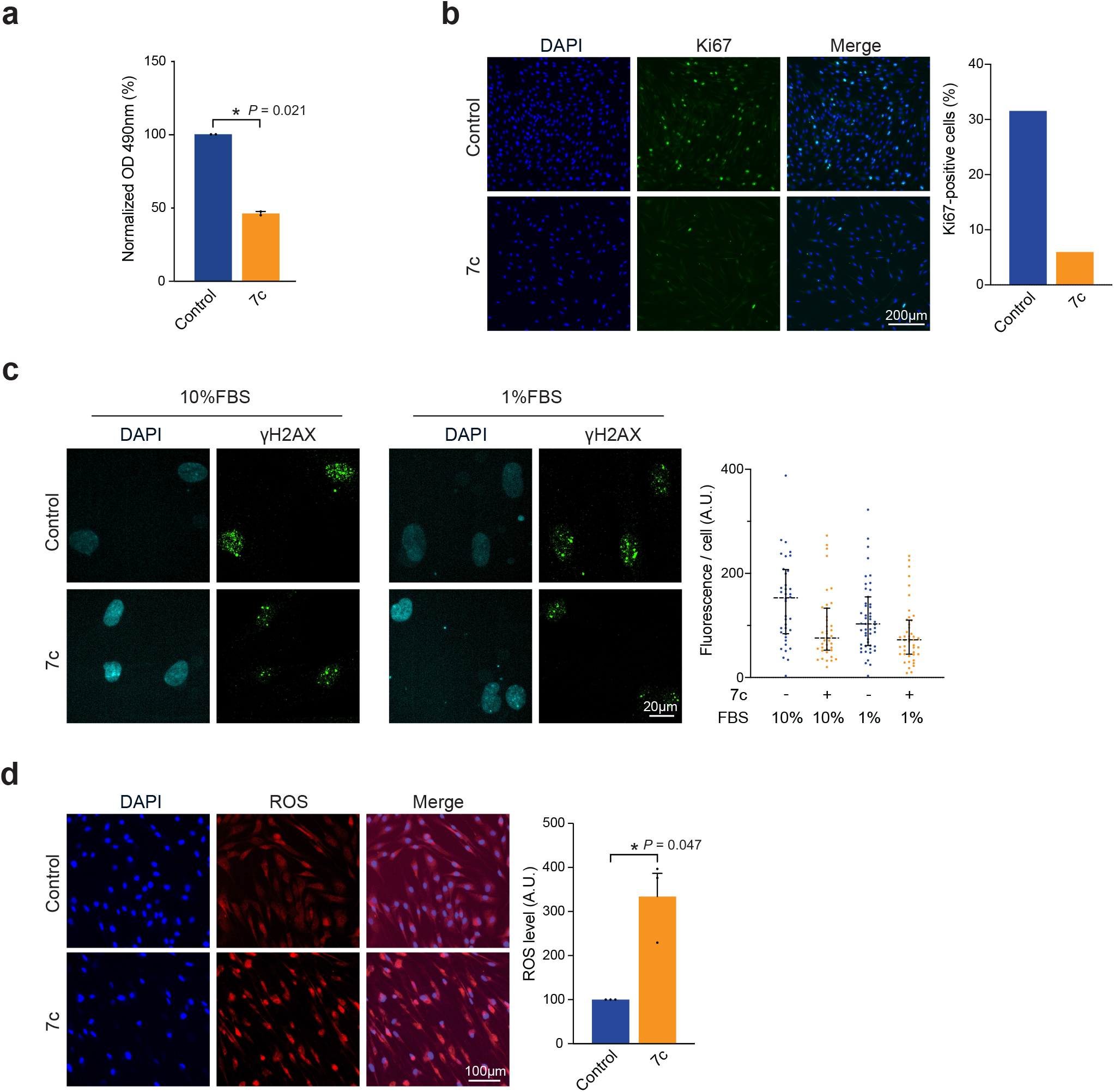
ciPR via 7 chemicals does not fully induce multiparameter rejuvenation. **a**, MTS quantification of cell density following 7c treatment until confluency. **b**, Immunofluorescence and quantification of Ki67 following 7c treatment (6 days, “6D”). **c**, Immunofluorescence and quantification of γH2AX following 7c treatment (6D) in proliferative (10%FBS) and non-proliferative (1%FBS) conditions (6D). **d**, Fluorescence detection and quantification of reactive oxygen species (ROS) following 7c treatment (6D). Data are mean ± SEM (**a, d**), median ± IQR (**c**). (**a-b**) n=2, (**c**) n=1, (**d**) n=3. Statistical significance was assessed by comparison to untreated control using paired two-tailed *t*-test (**a, d**). OD, Optical Density.

Next, in order to investigate whether the decrease in DNA damage upon 7c treatment was independent of cell cycle impairment, we tested the effect of 7c treatment under non-proliferative conditions in the presence of low-serum culture conditions (i.e. 1% FBS culture media). Remarkably, regardless of growth conditions and proliferation, 7c treatment still induced a reduction in γH2AX levels (Fig. 2c), suggesting that the impact of 7c on DNA damage is independent from its effects on proliferation. In addition, to gain further insight into the metabolic changes induced upon 7c treatment, we investigated the levels of reactive oxygen species (ROS), which are associated with mitochondrial function, cellular stress and DNA damage (Shields, Traa and Van Raamsdonk, 2021). Notably, a significant increase in reactive oxygen species (ROS) in aged fibroblasts was observed upon 7c treatment (Fig. 2d). Thus, although 7c treatment can decrease γH2AX levels, we find that it also leads to impaired proliferative capacity, even at low concentrations, and an upregulation of ROS. Taken together, chemical induced partial reprogramming via 7c treatment in aged human fibroblasts lacks the multiparameter rejuvenation associated with OSKM-induced reprogramming. In particular, 7c treatment results in the improvement of several key aging phenotypes such as DNA damage, epigenetic dysregulation, and senescence markers while at the same time leading to an impairment of proliferation, increased ROS, and upregulation of *IL6*. These results suggest that further optimization of chemical reprogramming may be required.

### A reduced reprogramming cocktail efficiently improves multiple molecular hallmarks of aging

In order to create an optimized cocktail for the amelioration of age-associated phenotypes by chemical reprogramming, we first sought to remove compounds with deleterious effects, while still retaining the three key functional categories of chemical reprogramming compounds: epigenetic modifiers, cell signaling modifiers, and metabolic switchers (Knyazer *et al*., 2021). First, using an MTS assay, we observed that while cell survival was unaffected or enhanced by Repsox or TCP treatment respectively, it was significantly impaired by CHIR99021, DZNep, Forskolin, TTNPB, and VPA, in agreement with previous publications and suggesting their removal (Fig. S2b) (Wu *et al*., 2006; Jung *et al*., 2008; Rodriguez *et al*., 2013; Fujiwara *et al*., 2014). In addition, DZNep has a known S-adenosylhomocysteine (SAH) hydrolase mediated inhibitory effect on the H3K27 methyltransferase EZH2, further supporting its exclusion (Girard *et al*., 2014). The remaining compounds, TCP and Repsox, met the selection criteria for chemical reprogramming functional categories and were thus selected. Therefore, we next treated aged human fibroblasts with this reduced two-chemical cocktail (2c) for 6 days to determine its effect on aging hallmarks. Strikingly, similar to our previous results with 7c, γH2AX levels were significantly decreased upon 2c treatment (Fig. 3a). Furthermore, improvement on yH2AX levels was observed when cells were treated with 2c following addition of the DNA damaging agent doxorubicin (Fig. S3b). Moreover, 2c significantly increased both H3K9me3 and H3K27me3 levels (Fig. 3b-c). Taken together, these data indicate that 2c treatment improves DNA damage and heterochromatin marks similar to 7c treatment.

**Figure 3.**
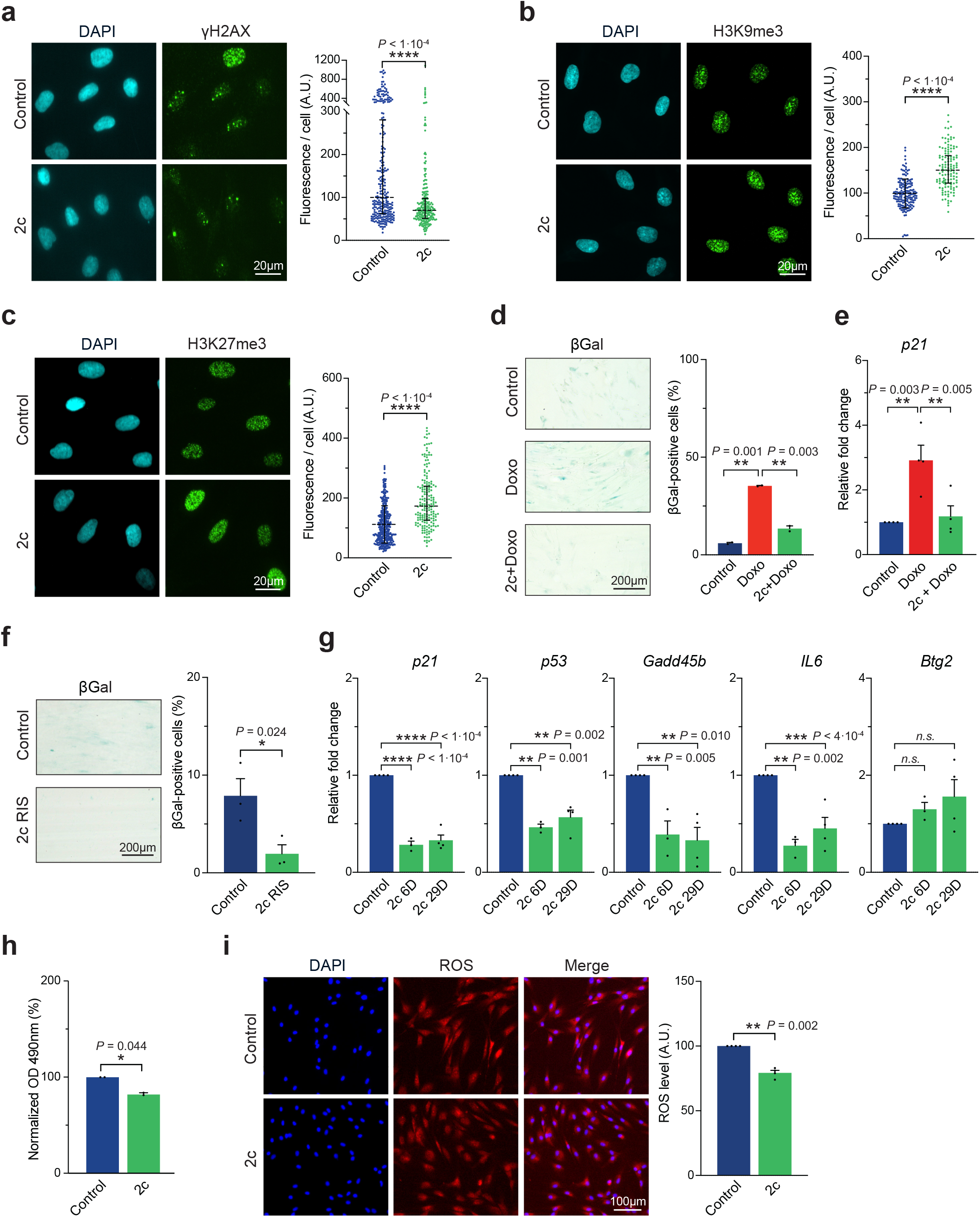
Optimized cocktail (2c) efficiently improves multiple molecular hallmarks of aging. **a**, Immunofluorescence and quantification of γH2AX following TCP + Repsox (2c, 5 µM each) treatment (6 days, “6D”). **b-c**, Immunofluorescence and quantification of H3K9me3 (**b**) and H3K27me3 (**c**) following 2c treatment (6D). **d**, Senescence-associated beta-galactosidase (SA-beta-gal) staining and quantification following Doxorubicin (100 nM) in 2c pre-treated fibroblasts (6D). **e**, mRNA levels of senescence-associated *p21* expression following Doxorubicin (100 nM) in 2c pretreated fibroblasts (29D). **f**, SA-beta-gal staining and quantification of replicative induced senescence (RIS) following long-term (29D) 2c treatment. **g**, mRNA levels of senescence-associated and age-related stress response genes in the *p53* tumor suppressor pathway following 2c treatment. **h**, MTS quantification of cell density following 2c treatment until confluency. **i**, Fluorescence detection and quantification of ROS following 2c treatment (6D). Data are median ± IQR (**a**), mean ± SEM (**b-i**). (**a-c, e-g, i**) n≥3, (**d, h**) n=2. Statistical significance was assessed by comparison to untreated control using paired two-tailed *t*-test (**f, h-i**), one-way ANOVA and Dunnett correction (**d-e, g**). OD, Optical Density.

Next, we sought to determine the impact of 2c treatment on cellular senescence in both a genotoxic stress induced senescence model using doxorubicin application and RIS after multiple passages. First, doxorubicin induced senescent cells pretreated with 2c showed a significant decrease in senescence-associated beta-galactosidase (SA-beta-gal) levels and *p21* gene expression compared to untreated controls (Fig. 3d-e). Interestingly, 2c significantly decreased SA-beta-gal levels only when added prior to induction, indicating a protective rather than senolytic effect upon genotoxic treatment (Fig. S3c). Moreover, in our RIS model, SA-beta-gal levels were significantly decreased in aged fibroblasts with continuous 2c treatment (Fig. 3f). In addition, senescence-associated and age-related stress response genes *p21, p53*, and *IL6*, were also downregulated upon 2c treatment after 6 days or 29 days of treatment (Fig. 3g). Taken together, these data show that 2c treatment reduces cellular senescence and significantly decreases *IL6* levels in contrast to 7c treatment.

Next, as 7c treatment previously led to impaired proliferation and increased ROS levels, we sought to determine the impact of 2c on these cellular phenotypes. Importantly, 2c treatment had only a mild effect on cell proliferation compared to the 50% decrease induced by 7c treatment (Fig. 3h). In addition, in clear contrast to the impact of 7c, 2c treatment significantly decreased ROS levels in aged fibroblasts, indicating that 2c can markedly improve cellular homeostasis (Fig. 3i). Similarly, we observed an improvement in ROS levels with 2c treatment when cells were co-treated with the pro-oxidant Antimycin A (Fig. S3a).

Overall, these results show that the reduced 2c cocktail is able to improve multiple age-related hallmarks including genomic instability, epigenetic dysregulation, and cellular senescence. Importantly, treatment with 2c had a minor effect on proliferation and led to a decrease in ROS levels. Taken together, these data indicate that 2c is an optimized cocktail capable of improving multiple age-related markers in vitro.

### 2c treatment increases *C. elegans* lifespan

Finally, in order to determine whether our 2c cocktail could also impact biological aging in vivo, we tested the effect of 2c treatment on the lifespan of a commonly used aging model organism, the nematode *Caenorhabditis elegans*. Towards this goal, we monitored survival in *C. elegans* treated with either 2c, Repsox, or TCP at three different concentrations (50, 100, or 200 μM) alongside a vehicle control. Strikingly, we observed that 2c treatment at 50 μM was sufficient to extend *C. elegans* median lifespan from 19 to 27 days, corresponding to a 42.1% increase relative to vehicle control (Fig. 4a, e). To a lesser extent, Repsox or TCP alone at 50 μM also increased *C. elegans* median lifespan to 25 days, a 31.6% increase over vehicle control (Fig. 4a, e). These results indicate that Repsox and TCP are each able to extend median lifespan in *C. elegans*, and when combined as part of the 2c cocktail, can lead to an even greater increase in median lifespan.

**Figure 4.**
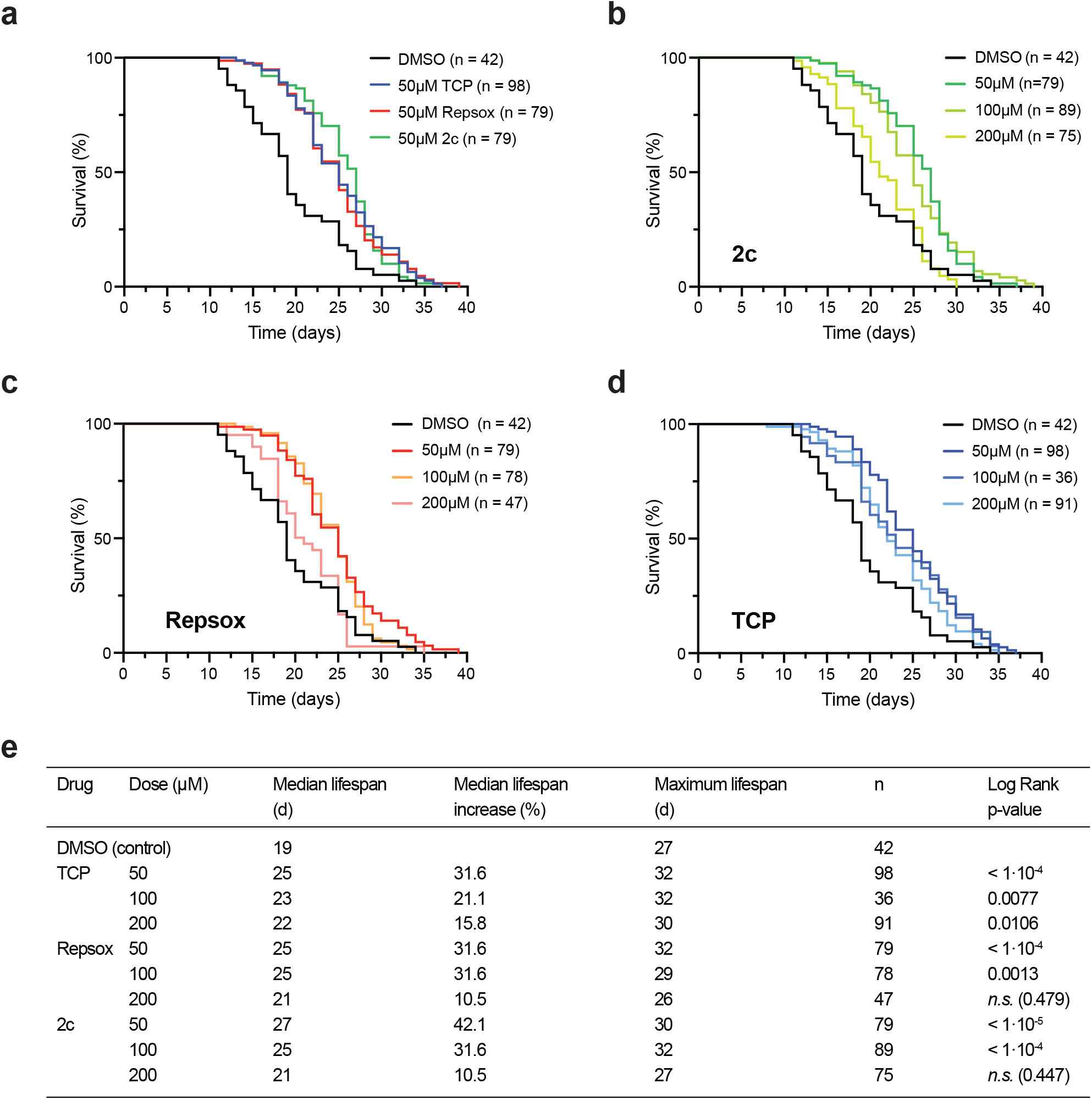
Treatment with 2c increases C. elegans lifespan. **a**, Survival of N2 *C. elegans* upon treatment of TCP (50 µM), Repsox (50 µM), and 2c (TCP + Repsox, 50 µM each). **b-d**, Survival of N2 *C. elegans* upon treatment with 2c (**b**), Repsox (**c**), and TCP (**d**) at 50, 100, or 200 µM. **e**, Summary of survival assay results including median lifespan, maximal (90%) lifespan, and statistical analyses. Median lifespan increase relative to vehicle control. Statistical significance was assessed by comparison to untreated control using Log-Rank (Mantel-Cox) test.

In addition, we observed dose-dependent effects across all treatments (Fig. 4b-e). Interestingly, the 2c cocktail or Repsox alone did not increase *C. elegans* lifespan at 200 μM (Fig. 4b-c, e), suggesting that this higher dose may impact off target mechanisms and be slightly toxic. On the other hand, TCP still increased median lifespan by 15.8% at 100 or 200 μM relative to vehicle controls even though it was most effective at 50 μM (Fig. 4d-e). Taken together, these data demonstrate that the optimized 2c cocktail can both ameliorate multiple aging hallmarks in aged human fibroblasts in vitro and extend *C. elegans* lifespan in vivo.

## DISCUSSION

The molecular identity and age of somatic cells have proven to be plastic states that can be reset by cellular reprogramming (Gurdon, 1962; Campbell *et al*., 1996; Takahashi and Yamanaka, 2006; Lapasset *et al*., 2011). As aging and age-associated diseases are a major societal burden, the need for aging interventions such as cellular reprogramming has grown (Mahmoudi, Xu and Brunet, 2019; Garmany, Yamada and Terzic, 2021). Although multiple groups have now demonstrated that in vivo partial reprogramming via transient application of OSKM can rejuvenate molecular hallmarks of aging, restore tissue function, and extend lifespan in mouse models, the risks of oncogenesis and inefficient gene delivery hinders clinical development (Ocampo *et al*., 2016; Kurita *et al*., 2018; de Lázaro *et al*., 2019; Lu *et al*., 2020; Rodríguez-Matellán *et al*., 2020; Sarkar *et al*., 2020; Chen *et al*., 2021; Cheng *et al*., 2022; Hishida *et al*., 2022). Interestingly, a more translational approach for the induction of cellular reprogramming based on the use of small molecules has been recently developed (Hou *et al*., 2013; Zhao *et al*., 2015; Cao *et al*., 2018; Guan *et al*., 2022). Still, the effects of small molecule induced cellular reprogramming on aging hallmarks and lifespan were, until now, unknown. Here, we demonstrated for the first time that partial chemical reprogramming induces multiparameter rejuvenation of key aging hallmarks including genomic instability, epigenetic dysregulation, and cellular senescence in vitro while simultaneously extending the lifespan of *C. elegans* in vivo. In particular, we demonstrated that the seven chemical reprogramming cocktail, defined by Hou *et al*., was able to improve age-associated DNA damage, epigenetic alterations, and induce a unique transcriptomic profile enriched for developmental processes in aged human fibroblasts in vitro. Our observations that 7c significantly impaired proliferation and increased ROS levels might contribute to the previously observed low efficiency of mouse iPSC induction (Hou *et al*., 2013). We further revealed that an optimized two-compound cocktail (2c) is sufficient to decrease the levels of the DNA damage marker γH2AX, increase H3K9me3 and H3K27me3, prevent both replicative and genotoxic induced senescence, and decrease oxidative stress. Most importantly, we found that our 2c cocktail applied in vivo was able to extend the median lifespan of *C. elegans* by 42.1%. Interestingly, the highest doses of 2c or single compounds were less effective to extend lifespan indicating potential off-target effects that may require further optimization.

Previous reports have demonstrated that in vivo treatment with a modified small molecule reprogramming cocktail similar to Hou *et al*. could enhance regeneration in the liver and heart, thus providing proof of principle that treatment with these reprogramming-associated chemicals could benefit tissue repair (Tang and Cheng, 2017; Huang *et al*., 2018). On the other hand, we have now shown that chemical reprogramming can improve multiple molecular hallmarks of aging similarly to OSKM-induced reprogramming, and extend the lifespan of *C. elegans*. Multiparameter rejuvenation across aging hallmarks is a defining trait of cellular reprogramming albeit future work is required to properly identify the mechanisms responsible for these benefits (Chondronasiou *et al*., 2022; Gill *et al*., 2022). In this line, attempts to induce multiparameter amelioration are now emerging as a strategy for rejuvenation even outside the field of cellular reprogramming. In this regard, Shaposhnikov *et al*. recently demonstrated a synergistic effect by targeting multiple aging hallmarks simultaneously, producing a significant increase in *Drosophila melanogaster* lifespan compared to single interventions (Shaposhnikov *et al*., 2022).

Importantly, several translational advantages highlight the potential use of chemical induced partial reprogramming for the amelioration of age-associated phenotypes, including the fact that small molecules are cell permeable and therefore easy to deliver. Furthermore, their effects can be modulated via dosage, and are transient and reversible, thus avoiding oncogenic pitfalls associated with transcription factor induction (Zhao, 2019). In this proof of principle study, our observations indicate that chemical reprogramming represents both a valuable opportunity for the development of future anti-aging interventions, along with the mechanistic understanding of the complex inter-relationships of aging hallmarks and their respective amelioration.

## METHODS

### Cell culture and maintenance

Human dermal fibroblasts were freshly extracted using Collagenase I (Sigma, C0130) and Dispase II (Sigma, D4693) and cultured in DMEM (Gibco, 11960085) containing non-essential amino acids (Gibco, 11140035), GlutaMax (Gibco, 35050061), Sodium Pyruvate (Gibco, 11360039) and 10% fetal bovine serum (FBS, Hyclone, SH30088.03) at 37°C in hypoxic conditions (3% O2). Subsequently, fibroblasts were passaged and cultured according to standard protocols. Aged donor samples were of 56 and 83 years of age.

### Immunofluorescence staining

Cells were washed with fresh PBS and then fixed with 4% paraformaldehyde (Roth, 0964.1) in PBS at room temperature (RT) for 15 minutes. After fixation, cells were washed 3 times, followed by a blocking and permeabilization step in 1% bovine serum albumin (Sigma, A9647-50G) in PBST (0.2% Triton X-100 in PBS) for 60 min (Roth, 3051.3). Cells were then incubated at 4°C overnight with appropriate primary antibody, washed in PBS, followed by secondary antibody incubation with DAPI staining at RT for 60 min. Coverslips were mounted using Fluoromount-G (Thermofisher, 00-4958-02), dried at RT in the dark for several hours, stored at 4°C until ready to image and -20°C for long-term.

### Immunofluorescence imaging

Confocal image acquisition was performed using the Ti2 Yokogawa CSU-W1 Spinning Disk (Nikon), using the 100X objective and with 15 z-sections of 0.3 μm intervals. Appropriate lasers were used (405 nm and 488 nm) with a typical laser intensity set to 5-10% transmission of the maximum intensity for methylated histones, Ki67 and ROS, and 30-40% for phosphorylated histones. Exposure time and binning were established separately to assure avoidance of signal saturation.

### Antibodies and compounds

Antibodies were provided from the following companies. Abcam: anti-H3K27me3 (ab192985); Cell Signaling: anti-H3K9me3 (13969), anti-γH2AX (9718), anti-Ki67 (15580); Bioconcept: anti-H3 (13969); Sigma: anti-β-Actin (A2228); Thermofisher: anti-Rabbit (A32790); Agilent: anti-Rabbit Immunoglobulins/HRP (P0448), anti-Mouse Immunoglobulins/HRP (P0447); Roth: DAPI (6843.1) Compounds were purchased from the following companies. Thermo Fisher: DHE (D11347); Cayman: Valproic Acid (13033), CHIR99021 (13122), Repsox (14794), Forskolin (11018), Doxorubicin (15007); Acros Organics: TCP (130472500); APExBIO: DZNep (A8182); Seleckchem: TTNPB (S4627); Roth: X-beta-Gal (2315.3)

### RNA analysis

Total RNA was extracted using Monarch Total RNA Miniprep Kit (New England Biolabs, T2010S) according to manufacturer’s instructions with DNase treatment (Qiagen, 79254) for 15 minutes (1:8 in DNase buffer). Total RNA concentrations were determined using the Qubit RNA BR Assay Kit (Thermofisher, Q10211). cDNA synthesis was performed by adding 4 μL of iScript™ gDNA Clear cDNA Synthesis (Biorad, 1725035BUN) to 500ng of RNA sample and run in a Thermocycler (Biorad, 1861086) with the following protocol: 5 min at 25°C for priming, 20 min at 46°C for reverse transcription, and 1 min at 95°C for enzyme inactivation. Final cDNA was diluted 1:5 using autoclaved water and stored at - 20°C. qRT-PCR was performed using SsoAdvanced SYBR Green Supermix (Bio-Rad, 1725272) in 384-well PCR plates (Thermofisher, AB1384) using the QuantStudio™ 12K Flex Real-time PCR System instrument (Thermofisher). Forward and reverse primers (1:1) were used at a final concentration of 5 µM with 1 µL of cDNA sample. Primer sequences are listed in Table S1.

### RNA sequencing, processing, analysis

RNA-Seq library preparation and sequencing was performed by Novogene (UK) Company Limited on an Illumina NovaSeq 6000 in 150 bp paired-end mode. Raw FASTQ files were assessed for quality, adapter content and duplication rates with FastQC. Reads were aligned to the Human genome (GRCh38) using the STAR aligner (v2.7.9a) (Dobin *et al*., 2013) with ‘--sjdbOverhang 100’. Number of reads per gene was quantified using the featureCounts function in the subread package (Liao, Smyth and Shi, 2013). Ensembl transcripts were mapped to gene symbols using the mapIds function in the AnnotationDbi package (Pagès *et al*., 2022) with the org.Hs.eg.db reference package (Carlson, 2019). Raw counts were normalized by library size and converted to counts per million (CPM) for downstream analysis. Dimensionality reduction was performed via Principal Component Analysis (PCA) using the R software. Differentially expressed genes (DEG) were computed by the limma R package (Ritchie *et al*., 2015), by fitting a linear model on each gene, with an adjusted p-value of 0.05. Gene set enrichment analysis (GSEA) for gene ontology (GO) (Ashburner *et al*., 2000) was performed using the clusterProfiler package (Wu *et al*., 2021); (Ashburner *et al*., 2000), from the list of DEG (with a valid Entrez ID) ranked by logFoldChange. We used org.HS.eg.db as a reference, selected Biological Processes (BP) only, and an adjusted p-value of 0.05 (Bonferroni). Pathways were ranked by Normalized Enrichment Score (NES). Z-Score were calculated for each gene to plot as a heatmap.

### MTS cell proliferation assay

Cell viability and proliferation assays were performed by Tetrazolium MTS assay. Control and treated cells were cultured for 1 day in 96-well plates then treated with small molecules for 3 consecutive days before incubation with 120 μL fresh media containing 20 μL of CellTiter 96® AQueous One Solution (Promega, G3580) for 1 to 4 hours at 37°C in a humidified, 5% CO2 atmosphere. The amount of product formed was measured by recording the absorbance at 490nm using a BioTek Epoch 2 microplate reader. Relative proportion of viable cells was determined as a relative reduction of the optical density (OD) compared to control OD.

### Crystal violet staining

Cells were cultured as previously described for the MTS cell proliferation assay. Culture media was then carefully removed from wells and cells were washed three consecutive times with room temperature PBS (Gibco, 21600069) followed by 45 minutes incubation with crystal violet (Roth, T123.2) solution (Crystal Violet 0.05%, Formaldehyde 0.4%, Methanol 1% in PBS 1X). Plates were then washed by careful immersion in tap water 2 times, drained upside down, and air dried. Finally, solubilization in 1% SDS was performed on an orbital shaker until no dense areas of coloration persisted and absorbance was measured at 570nm using a BioTek Epoch 2 microplate reader. Relative proportion of viable cells was determined as a relative reduction of the optical density (OD) compared to control OD.

### Senescence-associated β-galactosidase assay

Senescence-associated beta-galactosidase (SA-βgal) assay was performed as described in the literature (Debacq-Chainiaux *et al*., 2009). Briefly, light fixation was performed on cells plated on glass coverslips using a solution of 3% paraformaldehyde and 0.2% glutaraldehyde in PBS buffer for 5 minutes. Fixation solution was then removed, wells were washed several times and stained overnight at 37°C in a CO2-free incubator in a solution of 40 mM citric acid/Na phosphate buffer, 5 mM K4[Fe(CN)6]3H2O, 5 mM K3[Fe(CN)6], 150 mM sodium chloride, 2 mM magnesium chloride and 1 mg/mL X-gal (Roth, 2315.1) with a pH of 5.9-6.0. Finally, coverslips were stained with DAPI, followed by standard immunofluorescence protocol, images were taken using bright-field microscopy, and proportion of β-Gal-positive cells was then quantified.

### Reactive Oxygen Species assay

Mitochondrial reactive oxygen species (ROS) were measured using the superoxide indicator dihydroethdium (DHE). Briefly, first, cells were incubated in fresh FBS-free media containing 5 μM DHE and incubated at 37°C in a humidified, 5% CO2 atmosphere for 30 minutes. Following incubation, wells were washed with room temperature PBS, fixed with 4% paraformaldehyde for 15 min and then stained with DAPI. Immediately, images were taken at 554 nm and standard immunofluorescence imaging protocol was followed.

### Quantification and statistical analysis

Analysis of immunofluorescence microscopy images was performed using ImageJ. A minimum of 50-100 cells were imaged per condition. Maximal projections of z-stacks were analyzed and total fluorescence intensity per cell were determined.

All statistical parameters such as statistical analysis, statistical significance and n values are reported in the figure legends. Statistical analyses were performed using GraphPad Prism 9.0.0. Outliers were systematically removed using the ROUT method (Q=1%). For in vivo experiments, n corresponds to the numbers of animals and statistical analysis was performed using SPSS 27.0.1.0 (IBM® SPSS® Statistics).

### *C. elegans* strains and maintenance

Wildtype *C. elegans* (N2) were obtained from the Caenorhabditis Genetics Center (CGC), University of Minnesota, USA. N2 wildtype worms were maintained at 20°C and were grown on standard Nematode Growth Media (NGM) plates.

### Lifespan analysis

Animals were synchronized and lifespan analyses were conducted at 20°C as previously described (Porta-de-la-Riva *et al*., 2012) and were transferred onto NGM plates containing treatment or vehicle at stage L4. TCP was dissolved in water at 100 mM and Repsox was dissolved in DMSO at 200 mM. TCP and Repsox were added directly into the molten agar to a final concentration of 50, 100 or 200 μM each before pouring. After proper drying of plates, UV killed OP50 bacteria were seeded (150 μl of 120 mg/ml UV killed OP50 per P60 plate) and FUDR (150 μM) added as reproductive suppressant. Treated and control plates contained an equivalent DMSO concentration. Animals that crawled off the plate or displayed extruded internal organs were censored. Lifespan analyses were assessed manually by counting live and dead animals based on movement.

## AUTHOR CONTRIBUTIONS

A.O., P.T.P. and L.S. designed the study. A.O. and L.S. were involved in all experiments, data collection, analysis and interpretation. A.O., P.T.P. and L.S. prepared the figures and wrote the manuscript with input from all authors. L.S., G.B.P. and N.H.K. performed and analyzed in vivo experiments. C.M. performed skin fibroblast extractions. L.S. and K.P. prepared and analyzed RNA-seq data.

A.O. and P.T.P. provided assistance, supervision, and guidance.

## COMPETING INTERESTS

The authors declare no competing interests.

## ACKNOWLEDGMENTS

We would like to thank all members of the Ocampo laboratory for helpful discussions. In addition, we would like to thank the UNIL Cellular Imaging Facility and especially Luigi Bozzo for all technical training and guidance related to imaging.

## FUNDING

The study was supported by the Swiss National Science Foundation (SNSF) and the Canton Vaud.

---

**Figure S1.**
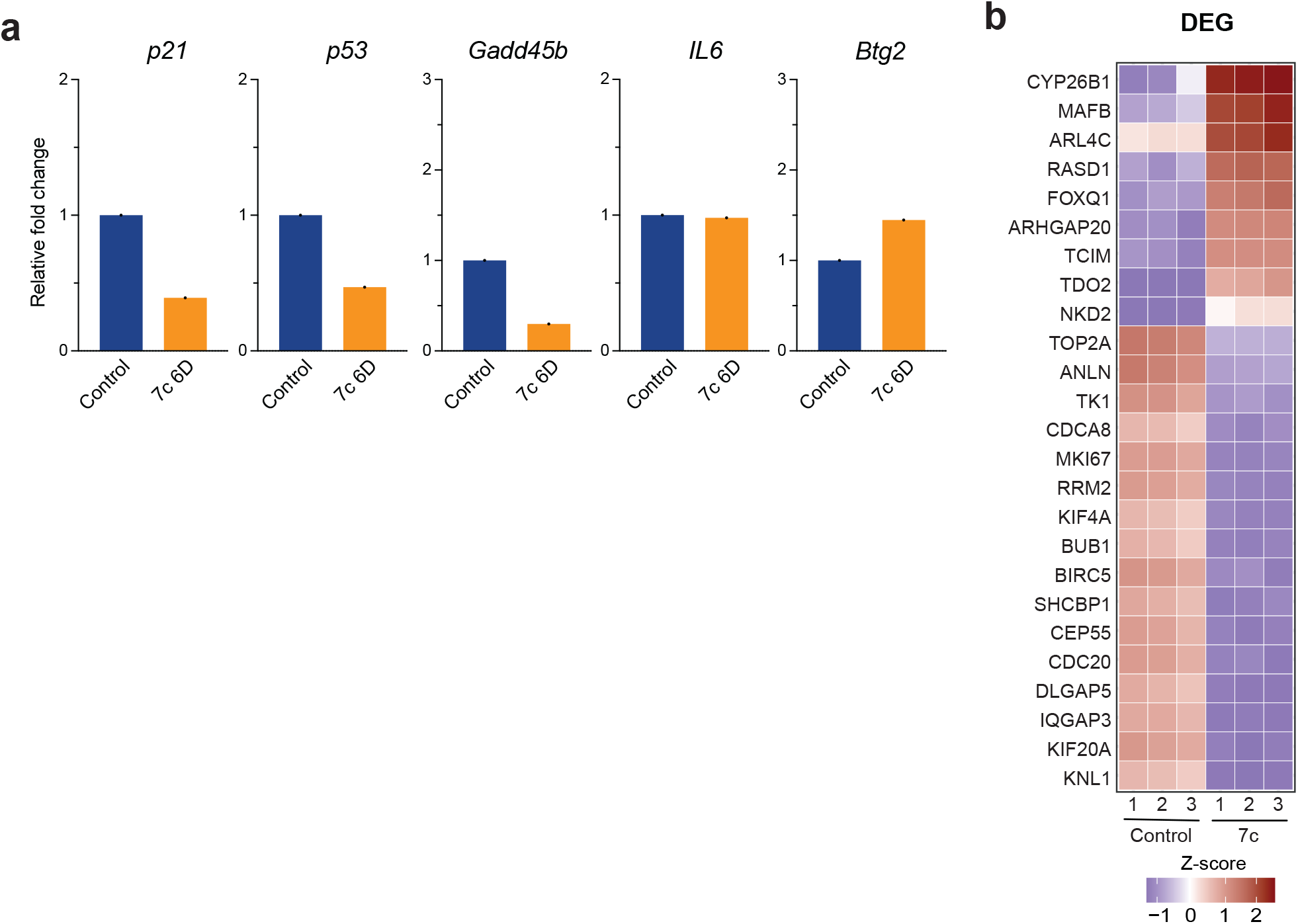
Gene expression analysis of chemical induced partial reprogramming with 7c treatment. **a**, mRNA levels of senescence-associated and age-related stress response genes in the *p53* tumor suppressor pathway following 7c treatment (6 days). **b**, Top differentially expressed genes following 7c treatment relative to untreated controls in human fibroblasts. (**a**) n=1, (**b**) n=3. DEG, differentially expressed genes.

**Figure S2.**
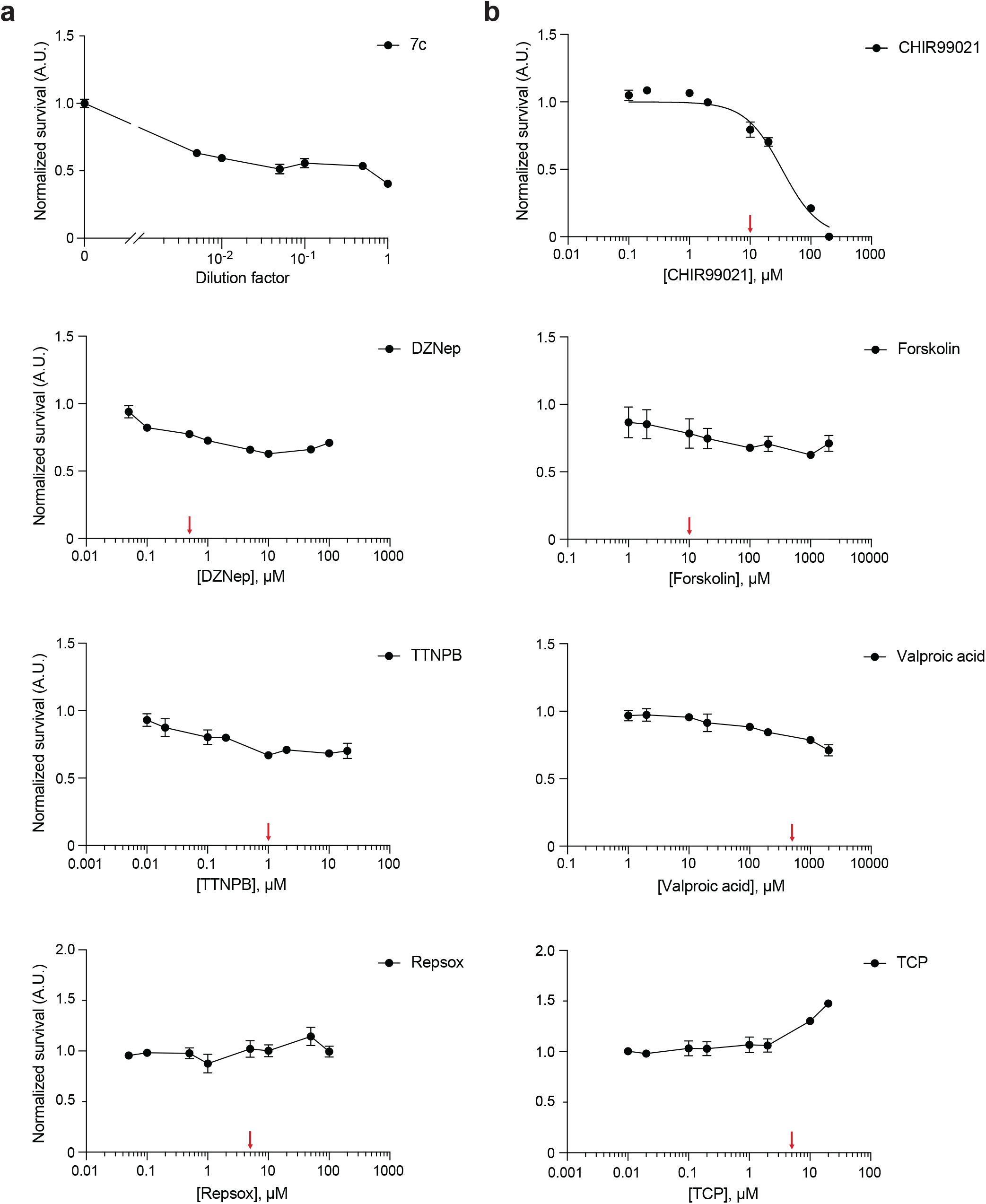
Serial dilution of the reprogramming chemicals. **a**, Crystal violet quantification of cell density following treatment until confluency with serial dilutions of the 7c reprogramming cocktail. **b**, MTS quantification of cell density following treatment until confluency with different concentration of the reprogramming chemicals. Red arrows indicate experimental concentrations. Initial concentrations noted in Table S2. Nonlinear regression displayed when possible. Data are mean ± SEM.

**Figure S3.**
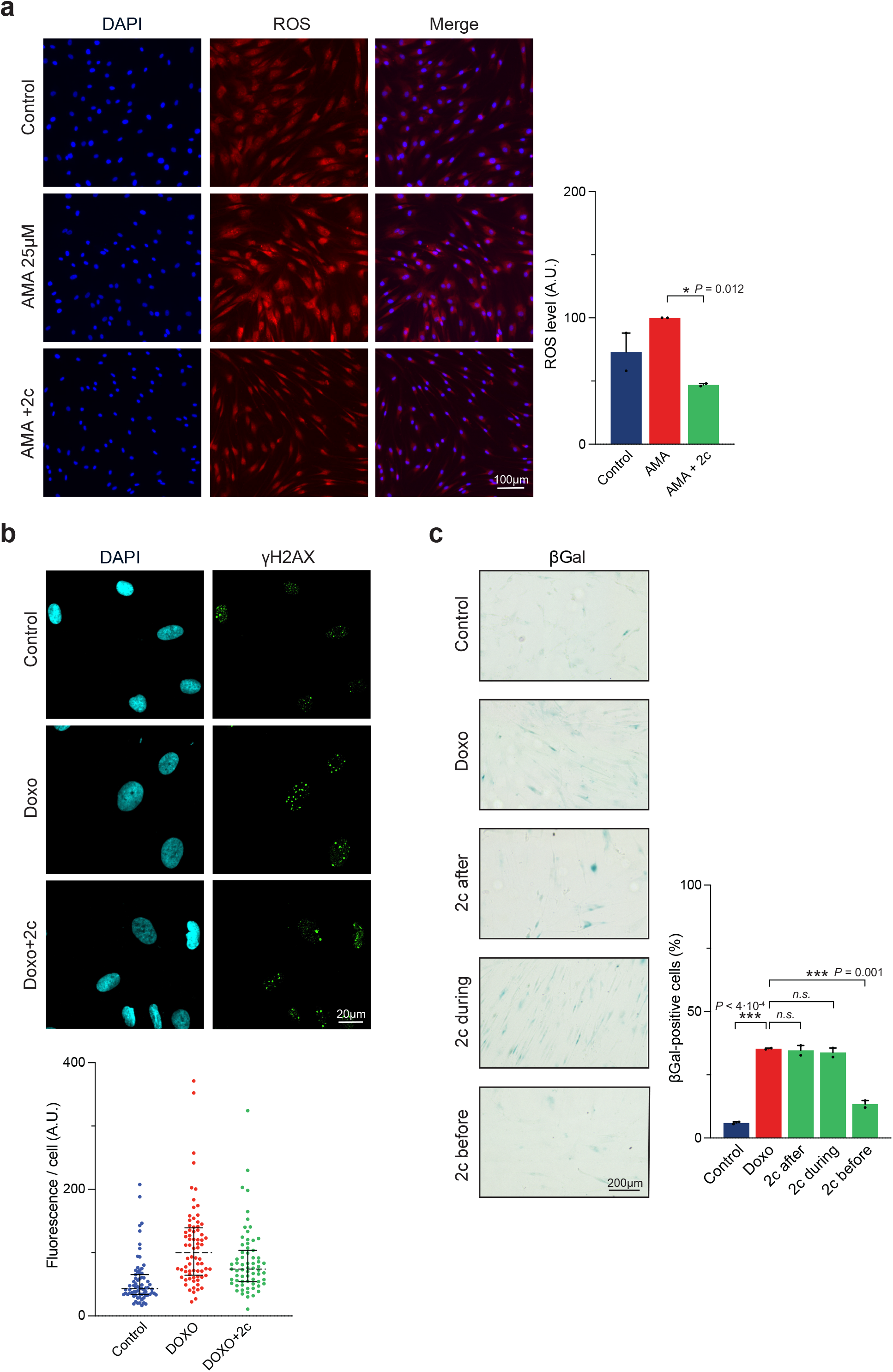
Reduced 2c cocktail efficiently ameliorates multiple hallmarks of aging. **a**, Fluorescence detection and quantification of ROS following AMA (25 µM) and 2c treatment (6 days, “6D”). **b**, Immunofluorescence and quantification of γH2AX following 2c treatment. **c**, Senescence-associated beta-galactosidase (SA-beta-gal) staining and quantification in 2c treated fibroblasts before, during, and after Doxorubicin (100 nM) treatment (6D). Data are mean ± SEM (**a, c**), median ± IQR (**b**). (**a-c**) n=2, (**b**) n=1. Statistical significance was assessed by comparison to untreated control using paired two-tailed *t*-test (**a**), one-way ANOVA and Dunnett correction (**c**).

**Table S1.**
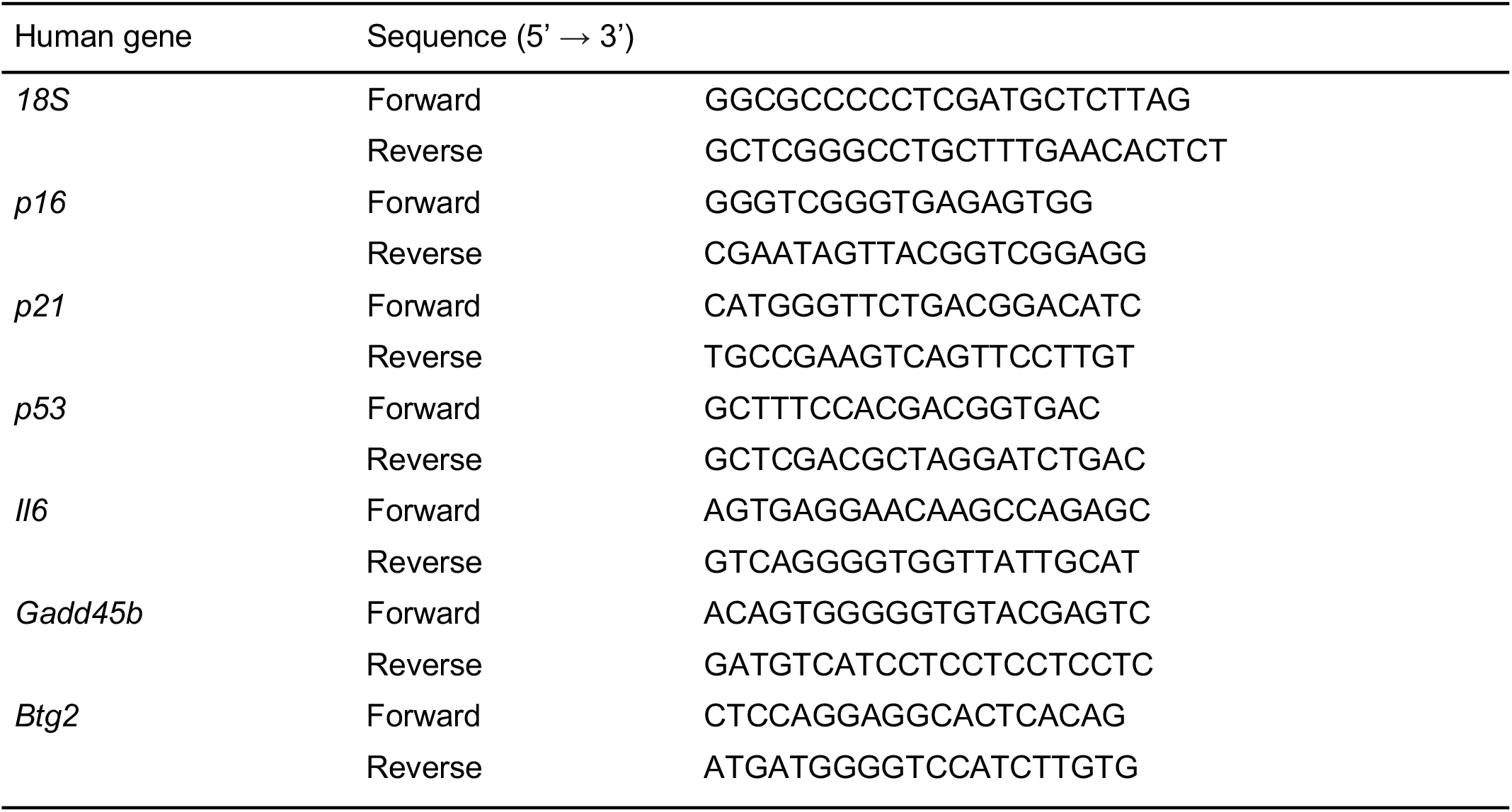
Primers set for qRT-PCR.

**Table S2.**
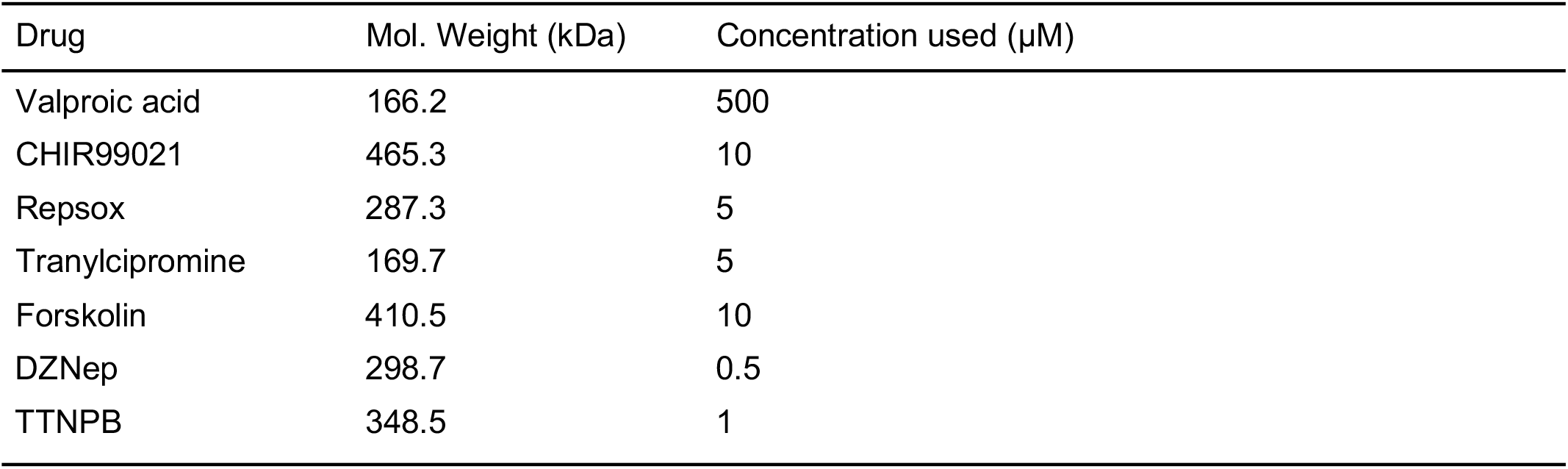
Table of reprogramming chemicals and respective concentrations used.

## Notes

### Competing Interest Statement

The authors have declared no competing interest.

